## Supplemental Information for "Cluster Analysis of Medicinal Plants and Targets Based on Multipartite Network"

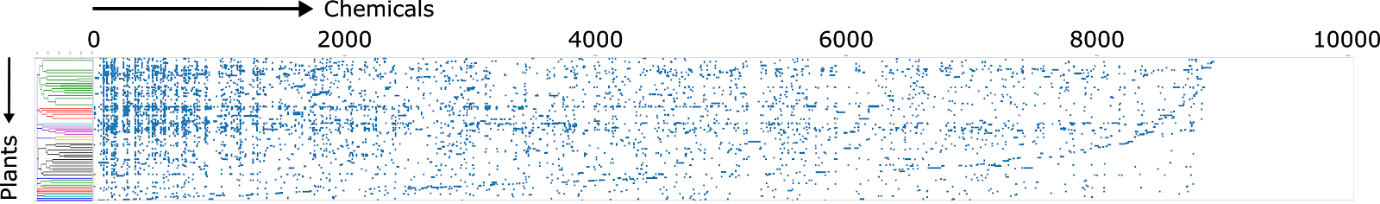

**Figure S1.** Visualization of non-zero values of the data matrix for the plants and chemicals, where the rows correspond to the plants and the columns correspond to the chemicals. The rows were permuted according to the order determined by the hierarchical clustering of the plants.

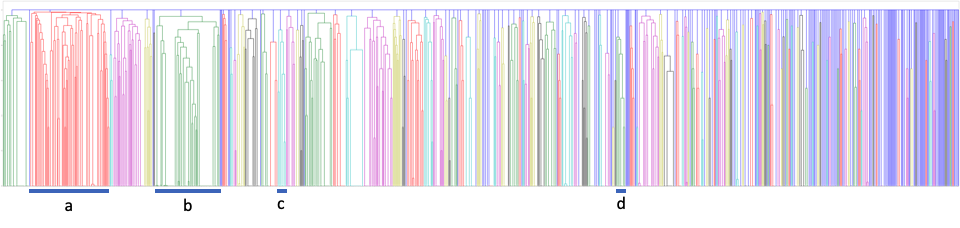

**Figure S2.** The whole dendrogram for clustering of plants obtained by the hierarchical clustering analysis. **a-d**: the dendrograms of **Figures 2A, 2C**, **2E** and **2H**, respectively.

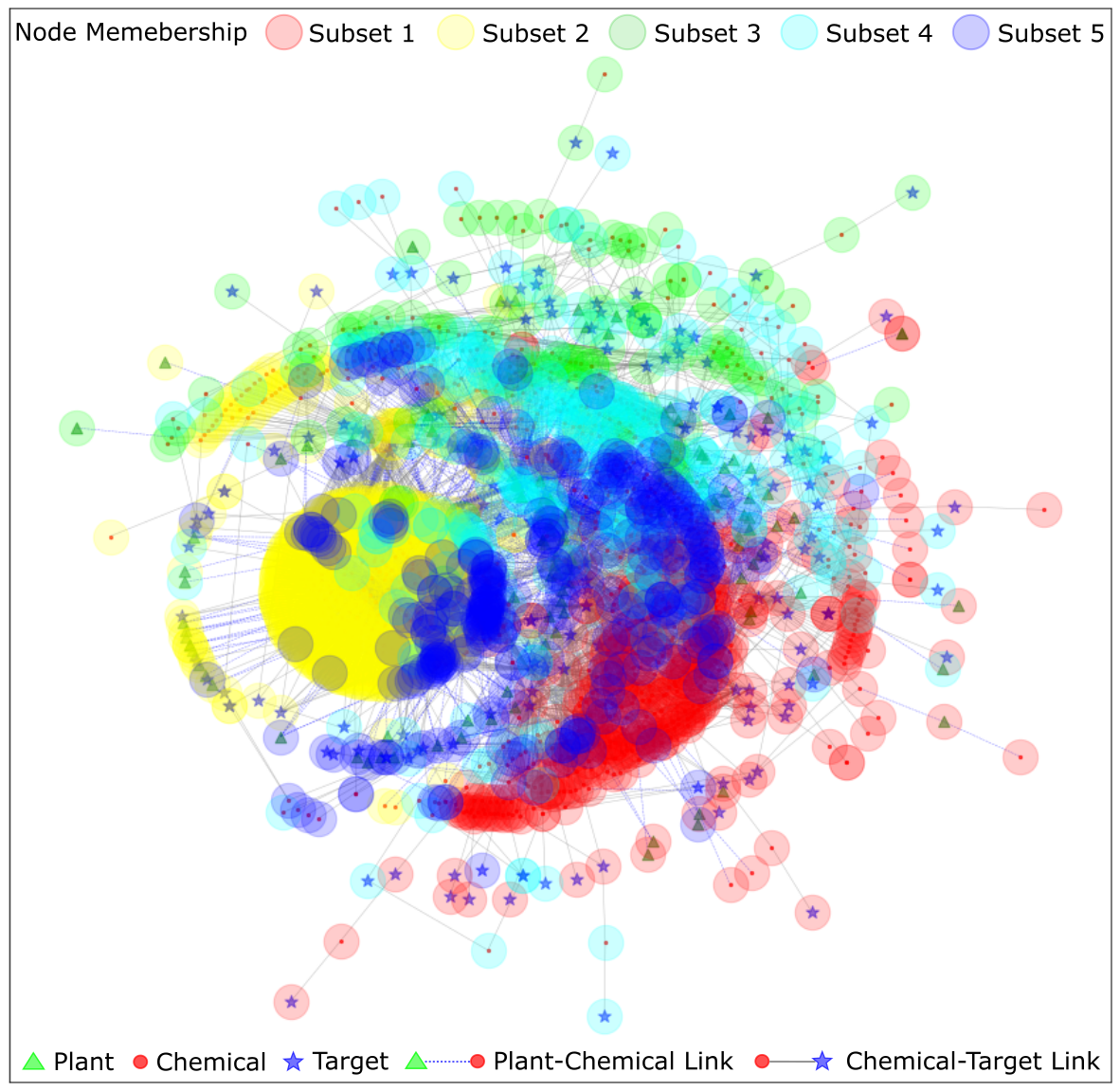

(A)

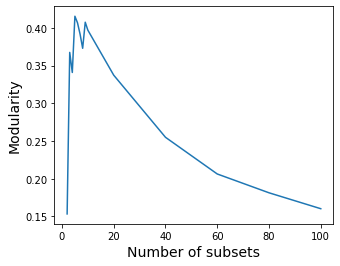

(B)

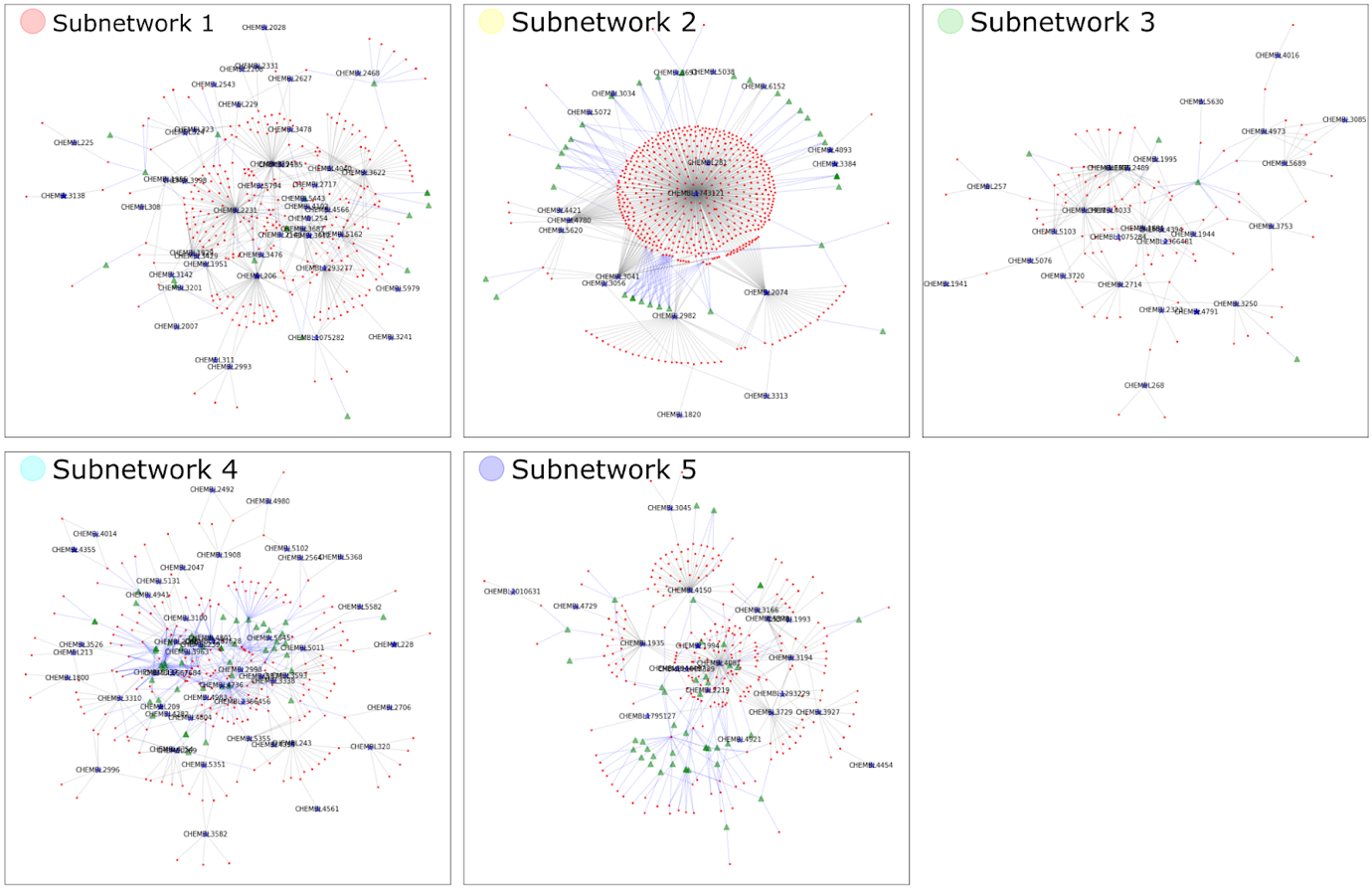

(C)

**Figure S3.** Visualization of the results of the spectral co-clustering. **(A)** Spectral co-clustering of the whole network. **(B)** The modularity values for the determination of the number of clusters for the spectral co-clustering algorithm. **(C)** The subnetworks induced by each of the five subsets of nodes obtained by the spectral co-clustering. The 10% of the whole nodes were randomly selected for visualization.

**Table S1**. The structural properties of the whole network and the five subnetworks obtained by the spectral co-clustering algorithm.

| Network | Number of Nodes | | | | Number of Edges | | | Diameter | Average Bipartite Clustering Coefficient | Degree Assortativity |
| --- | --- | --- | --- | --- | --- | --- | --- | --- | --- | --- |
|  | Np | Nc | Nt | N | Epc | Ect | E |  |  |  |
| Whole | 1138 | 10051 | 1224 | 12413 | 34549 | 100290 | 134839 | 5 | 0.0946 | -0.1954 |
| Sub 1 | 276 | 3459 | 124 | 3859 | 3915 | 29493 | 33408 | 4 | 0.2116 | -0.3936 |
| Sub 2 | 327 | 2538 | 136 | 3001 | 3946 | 13456 | 17402 | 5 | 0.1595 | -0.3672 |
| Sub 3 | 34 | 676 | 177 | 887 | 205 | 5491 | 5696 | 5 | 0.1544 | -0.4156 |
| Sub 4 | 141 | 1907 | 361 | 2409 | 1564 | 14118 | 15682 | 4 | 0.1315 | -0.3814 |
| Sub 5 | 360 | 1471 | 426 | 2257 | 9751 | 5893 | 15644 | 5 | 0.1436 | -0.2290 |

Np=number of plant nodes, Nc=number of chemical nodes, Nt=number of target nodes, N=total number of nodes, Epc=number of plant-chemical edges, Ect=number of chemical-target edges, E=total number of edges.

**Table S2**. List of plants illustrated in **Figures 2A** and **2B**.

| **Plant ID** | **Scientific Name** | **Latin Name** | **Family** |
| --- | --- | --- | --- |
| **M0000066** | *Euphorbia kansui* | Euphorbiae Kansui Radix | Euphorbiaceae |
| **M0000069** | *Citrus suavissima* | Citri Suavissimae Exocarpium | Rutaceae |
| **M0000072** | *Glycyrrhiza uralensis* | Glycyrrhizae Praeparatus Pulvis | Leguminosae |
| **M0000075** | *Glycyrrhiza inflata* | Glycyrrhizae Radix | Leguminosae |
| **M0000079** | *Glycyrrhiza inflata* | Glycyrrhizae Resina | Leguminosae |
| **M0000080** | *Glycyrrhiza uralensis* | Glycyrrhizae Radicii Virgula | Leguminosae |
| **M0000088** | *Pinellia ternata* | Pinelliae Praeparatum cum Zingiberis | Araceae |
| **M0000110** | *Dianthus superbus* | Dianthi Herba | Caryophyllaceae |
| **M0000127** | *Zizyphus jujuba var. inermis* | Zizyphi Fructus | Rhamnaceae |
| **M0000216** | *Melia azedarach* | Meliae Cortex | Meliaceae |
| **M0000222** | *Melia azedarach* | Meliae Fructus | Meliaceae |
| **M0000224** | *Melia azed-arach var. japonica* | Meliae Cortex | Meliaceae |
| **M0000262** | *Cucumis melo* | Melo Pediculus | Cucurbitaceae |
| **M0000264** | *Sorghum vulgare* | Sorghi Acetum | Gramineae |
| **M0000265** | *Hordeum vulgare* | Hordei Acetum | Gramineae |
| **M0000334** | *Hordeum vulgare* | Hordei Fructus | Gramineae |
| **M0000567** | *Raphanus sativus* | Raphani Radix | Cruciferae |
| **M0000581** | *Kochia scoparia* | Kochiae Fructus | Chenopodiaceae |
| **M0000607** | *Strobilenthes flaccidifolia* | Strobilenthis Sedimentum | Acanthaceae |
| **M0000646** | *Raphanus sativus* | Raphani Semen | Cruciferae |
| **M0000771** | *Echinops latifolius* | Rhapontici Radix | Compositae |
| **M0000816** | *Lophatherum gracile* | Lophatheri Herba | Gramineae |
| **M0000854** | *Hordeum vulgare* | Hordei Semen | Gramineae |
| **M0000855** | *Hordeum vulgare* | Hordei Semen Pulvis | Gramineae |
| **M0000856** | *Hordeum vulgare* | Hordei Novella | Gramineae |
| **M0000877** | *Zizyphus vulgaris var. spinosus* | Jujubae Fructus | Rhamnaceae |
| **M0000878** | *Zizyphus jujuba var. inermis* | Jujubae Fructus | Rhamnaceae |
| **M0000990** | *Eucommia ulmoides* | Eucommiae Cortex | Eucommiaceae |
| **M0001001** | *Juncus effusus* | Junci Medulla | Juncaceae |
| **M0001022** | *Aristolochia contorta* | Aristolochiae Fructus | Aristolochiaceae |
| **M0001046** | *Calvatia gigantea* | Lasiosphaera Seu Calvatia | Rosacean |
| **M0001069** | *Portulaca oleracea* | Portulacea Herba | Portulacaceae |
| **M0001089** | *Vitex trifolia* | Viticis Fructus | Verbenaceae |
| **M0001108** | *Liriope platyphylla* | Liriopes Radix | Liliaceae |
| **M0001114** | *Hordeum vulgare var. hexastichon* | Hordei Fructus Germiniatus | Gramineae |
| **M0001115** | *Hordeum vulgare* | Hordei Fructus Germinatus | Gramineae |
| **M0001189** | *Aralia elata* | Araliae Cortex Lignum seu Radicis | Araliaceae |
| **M0001234** | *Ficus carica* | Fici Fructus | Moraceae |
| **M0001303** | *Pinellia pedatisecta* | Pinelliae Rhizoma | Araceae |
| **M0001305** | *Pinellia ternata* | Pinelliae Rhizoma Fermentata | Araceae |
| **M0001309** | *Eleocharis dulcis* | Eleocharitis Tuber | Cyperaceae |
| **M0001357** | *vulgaris Lamarck var. spinosus* | Zizyphi Spina | Rhamnaceae |
| **M0001377** | *Ampelopsis japonica* | Ampelopsis Radix | Vitaceae |
| **M0001523** | *Ficus pumila* | Fici Pumilae Caulis | Moraceae |
| **M0001541** | *Psoralea corylifolia* | Psoraleae Fructus | Leguminosae |
| **M0001581** | *Ailanthus altissima\|Ailanthus glandulosa* | Ailanthi Herba | Simaroubaceae |
| **M0001628** | *Torreya nucifera* | Torreyae Semen | Taxaceae |
| **M0001629** | *Torreya grandis* | Torreyae Semen | Taxaceae |
| **M0001663** | *Adenophora triphylla var. japonica* | Adenophorae Radix | Campanulaceae |
| **M0001710** | *Sambucus javanica* | Sambuci Radix | Caprifoliaceae |
| **M0001780** | *Gardenia jasminoides* | Gardeniae Fructus | Rubiaceae |
| **M0001808** | *Morus alba* | Mori Cortex | Moraceae |
| **M0001829** | *Cynanchum atratum* | Cynanchi Radix | Asclepiadaceae |
| **M0001836** | *Morus alba* | Mori Favilla | Moraceae |
| **M0001839** | *Morus alba* | Mori Fructus | Moraceae |
| **M0001855** | *Morus alba* | Mori Ramulus | Moraceae |
| **M0001940** | *Dendrobium loddigesii* | Dendrobii Herba | Orchidaceae |
| **M0002165** | *Cynomorium songaricum* | Cynomorii Herba | Cynomoriaceae |
| **M0002301** | *Centipeda minima* | Centipedae Herba | Compositae |
| **M0002470** | *Hordeum vulgare* | Hordei Germinatus | Gramineae |
| **M0002552** | *Mirabilis jalapa* | Fucus cum Jalapae Tuber | Nyctaginaceae |
| **M0002555** | *Mirabilis jalapa* | Fucus cum Jalapae Embryo | Nyctaginaceae |
| **M0002598** | *Acanthopanax sessiliflorus* | Acanthopanacis Cortex | Araliaceae |
| **M0002661** | *Lindera aggregata* | Linderae Radix | Lauraceae |
| **M0002913** | *Citrus suavissima* | Citri Suavissimae Fructus | Rutaceae |
| **M0002915** | *Artemisia anomala\|Artemisia viridissima* | Artemisiae Anomalae Herba | Compositae |
| **M0002923** | *Ulmus macrocarpa* | Ulmi Cortex | Ulmaceae |
| **M0002991** | *Coix lachryma-jobi var. ma-yeun* | Coicis Radix | Gramineae |
| **M0002992** | *Coix lachryma-jobi* | Coicis Radix | Gramineae |
| **M0002993** | *Coix lachryma-jobi var. ma-yeun* | Coicis Semen | Gramineae |
| **M0002994** | *Coix lachryma-jobi* | Coicis Semen | Gramineae |
| **M0003064** | *Glycyrrhiza uralensis* | Glycyrrhizae Radix Praeparata | Leguminosae |
| **M0003088** | *Spirodela polyrrhyza* | Spirodelae Herba | Lemnaceae |
| **M0003284** | *Tamarix chinensis* | Tamaricis Ramulus | Tamaricaceae |
| **M0003373** | *Adenophora trachelioides* | Trachelioidis Radix | Campanulaceae |
| **M0003415** | *Zizyphus jujuba var. inermis* | Zizyphi Radix | Rhamnaceae |
| **M0003417** | *Zizyphus jujuba var. inermis* | Zizyphi Folium | Rhamnaceae |
| **M0003492** | *Kochia scoparia* | Kochiae Herba | Chenopodiaceae |
| **M0003630** | *Euphorbia lathyris* | Euphorbiae Lathyridis Semen | Euphorbiaceae |
| **M0003670** | *Aristolochia contorta* | Aristolochiae Herba | Aristolochiaceae |
| **M0003719** | *Cucumis melo* | Melo Fructus | Cucurbitaceae |
| **M0003720** | *Cucumis melo* | Melo Semen | Cucurbitaceae |
| **M0003721** | *Cucumis melo* | Melonis Pedicelus | Cucurbitaceae |
| **M0003731** | *Baphicacanthus cusia* | Indigo Pulverata Levis | Acanthaceae |
| **M0003752** | *Morus alba* | Mori Radicis Cortex | Moraceae |
| **M0003786** | *Celosia argentea* | Celosiae Semen | Amaranthaceae |
| **M0003830** | *Sorghum vulgare* | Sorghi Radix | Gramineae |
| **M0003831** | *Dichroa febrifuga\|Orixa japonica* | Dichroae Arbor Novella | Saxifragaceae |
| **M0003858** | *Sorghum vulgare* | Sorghi Semen | Gramineae |
| **M0003887** | *Gardenia jasminoides* | Gardeniae Semen | Rubiaceae |
| **M0003969** | *Psoralea corylifolia* | Psoraleae Semen | Leguminosae |
| **M0004009** | *Patrinia scabiosaefolia* | Patriniae Radix | Valerianaceae |
| **M0004073** | *Prunella vulgaris* | Prunellae Spica | Labiatae |
| **M0004247** | *Cucumis sativus* | Cucumeris Fructus | Cucurbitaceae |
| **M0004398** | *Cudrania tricuspidata* | Cudraniae Folium | Moraceae |

**Table S3**. List of plants illustrated in **Figures 2C** and **2D**.

| **Plant ID** | **Scientific Name** | **Latin Name** | **Family** |
| --- | --- | --- | --- |
| **M0000060** | *Saccharum sinensis* | Sacchari Tuber | Gramineae |
| **M0000124** | *Caesalpinia sappan\|Cercis chinensis* | Fucus cum Caesalpiniae | Leguminosae |
| **M0000179** | *Areca catechu* | Arecae Semen | Palmae |
| **M0000227** | *Cannabis sativa* | Vetus Cannabis Solea | Cannabinaceae |
| **M0000263** | *Oryza sativa* | Oryzae Acetum | Gramineae |
| **M0000286** | *Oryza sativa* | Oryzae Fructus Germinatus | Gramineae |
| **M0000557** | *Oryza sativa* | Oryzae Glutinosa Medulla | Gramineae |
| **M0000559** | *Oryza sativa* | Oryzae Radix seu Rhizoma | Gramineae |
| **M0000564** | *Oryza sativa* | Oryzae Semen | Gramineae |
| **M0000565** | *Oryza sativa* | Alcohol cum Oryzae Semen | Gramineae |
| **M0000803** | *Santalum album* | Santali Albae Lignum | Santalaceae |
| **M0000821** | *Angelica sinensis* | Angelicae Gigantis Radix | Umbelliferae |
| **M0000842** | *Glycine max* | Glycinis Semen | Leguminosae |
| **M0000844** | *Glycine max* | Glycinis Semen Germinatum | Leguminosae |
| **M0000846** | *Glycine max* | Glycine Semen Germinatum | Leguminosae |
| **M0000850** | *Cannabis sativa* | Cannabis Herba | Cannabinaceae |
| **M0000852** | *Cannabis sativa* | Cannabis Semen | Cannabinaceae |
| **M0000862** | *Areca catechu* | Arecae Pericarpium | Palmae |
| **M0000864** | *Allium sativum* | Allii Bulbus | Liliaceae |
| **M0000913** | *Oryza sativa* | Oryzae Semen Germinatus | Gramineae |
| **M0000938** | *Benincasa cerifera\|Benincasa hispida* | Benincasae Fructus | Cucurbitaceae |
| **M0000939** | *Benincasa cerifera\|Benincasa hispida* | Benincasae Caulis | Cucurbitaceae |
| **M0000940** | *Benincasa cerifera\|Benincasa hispida* | Benincasae Folium | Cucurbitaceae |
| **M0000984** | *Glycine max* | Alcohol cum Glycinis Semen | Leguminosae |
| **M0000986** | *Glycine max* | Glycinis Semen Ptisanari | Leguminosae |
| **M0000992** | *Glycine max* | Glycinis Semen | Leguminosae |
| **M0001013** | *Agaricus campestris* | Agaricus Polyporus | Agaricaceae |
| **M0001014** | *Cannabis sativa* | Cannabis Radix | Cannabinaceae |
| **M0001058** | *Cannabis sativa* | Cannabis Folium | Cannabinaceae |
| **M0001062** | *Cannabis sativa* | Cannabis Fructus | Cannabinaceae |
| **M0001191** | *Auricularia auricula-judae* | Auriculariae Polyporus | Auriculariaceae |
| **M0001197** | *Auricularia auricula-judae* | Auriculariae Polyporus | Auriculariaceae |
| **M0001359** | *Santalum album* | Santali Albi Lignum | Santalaceae |
| **M0001418** | *Oryza sativa* | Oryzae Semen Pulvus | Gramineae |
| **M0001422** | *Saccharum sinensis* | Saccharum Alba | Gramineae |
| **M0001473** | *Oryza sativa* | Oryzae Ptisanari | Gramineae |
| **M0001540** | *Oryza sativa* | Alcohol Orizae cum Medico Fermento | Gramineae |
| **M0001639** | *Saccharum sinensis* | Saccharum Glacialis | Gramineae |
| **M0001659** | *Saccharum sinensis* | Saccharum | Gramineae |
| **M0001901** | *Scutellaria baicalensis* | Scutellariae Radix | Labiatae |
| **M0002086** | *Caesalpinia sappan\|Cercis chinensis* | Sappan Lignum | Anacardiaceae |
| **M0002150** | *Tricholoma matsutake* | Tricholomae Polyporus | Tricholomataceae |
| **M0002185** | *Polygonum hydropiper* | Polygoni Hydropiper Herba | Polygonaceae |
| **M0002221** | *Benincasa cerifera\|Benincasa hispida* | Benincasae Fructus | Cucurbitaceae |
| **M0002476** | *Glycine max* | Glycinis Semen | Leguminosae |
| **M0002477** | *Glycine max* | Glycinis Testa | Leguminosae |
| **M0002568** | *Oryza sativa* | Vinum Ferri | Gramineae |
| **M0002573** | *Glycine max* | Faba cum Sale Fermentoque Praeparata | Leguminosae |
| **M0002721** | *Pisum sativum* | Pisi Flos | Leguminosae |
| **M0002742** | *Polygonum hydropiper* | Polygoni Fructus | Polygonaceae |
| **M0002806** | *Cyathula capitata* | Achyranthis Radix | Amaranthaceae |
| **M0002807** | *Cyathula officinalis* | Achyranthis Radix | Amaranthaceae |
| **M0002810** | *Achyranthes bidentata* | Achyranthis Radix | Amaranthaceae |
| **M0002870** | *Glycine max* | Glycine Semen Nigrae | Leguminosae |
| **M0003033** | *Panax ginseng* | Ginseng Radix | Araliaceae |
| **M0003035** | *Panax ginseng* | Ginseng Nodus | Araliaceae |
| **M0003066** | *Sagittaria sagittifolia* | Sagittariae Tuber | Alismataceae |
| **M0003235** | *Oryza sativa* | Oryzae Testa | Gramineae |
| **M0003380** | *Capsella bursa-pastoris* | Capsellae Herba | Cruciferae |
| **M0003381** | *Capsella bursa-pastoris* | Capsellae Semen | Cruciferae |
| **M0003559** | *Oryza sativa* | Arida Oryza Distillata | Gramineae |
| **M0003622** | *Angelica sinensis* | Angelicae Sinensis Radix | Umbelliferae |
| **M0003764** | *Glycine max* | Liquor Salsus ex Faba | Leguminosae |
| **M0003765** | *Oryza sativa* | Purum Vinum Oryzae | Gramineae |
| **M0003853** | *Oryza sativa* | Vinum in Primo Vere | Gramineae |
| **M0003864** | *Oryza sativa* | Arida Oryza Praeparatum | Gramineae |
| **M0004061** | *Ricinus communis* | Ricini Folium | Euphorbiaceae |
| **M0004062** | *Ricinus communis* | Ricini Semen | Euphorbiaceae |
| **M0004267** | *Coriandrum sativum* | Coriandri Herba cum Radix | Umbelliferae |
| **M0004268** | *Coriandrum sativum* | Coriandri Herba cum Radix | Umbelliferae |
| **M0004269** | *Coriandrum sativum* | Coriandri Fructus | Umbelliferae |
| **M0004270** | *Glycine max* | Glycinis Semen Praeparatum | Leguminosae |
| **M0004286** | *Piper nigrum* | Pireris Nigri Fructus | Piperaceae |
| **M0004303** | *Alpinia galanga* | Galangae Fructus | Zingiberaceae |
| **M0004460** | *Glycine max* | Glycinis Semen Nigra | Leguminosae |
| **M0004463** | *Saccharum sinensis* | Saccharum Nigrum | Gramineae |
| **M0004468** | *Glycine max* | Glycinis Nigra Testa | Leguminosae |

**Table S4**. List of plants illustrated in  **Figures 2E** and **2F**.

| **Plant ID** | **Scientific Name** | **Latin Name** | **Family** |
| --- | --- | --- | --- |
| **M0000806** | *Phellodendron amurense* | Phellodendri Caulis | Rutaceae |
| **M0002558** | *Corydalis ternata* | Corydalis (Tuber) Rhizoma | Papaveraceae |
| **M0003700** | *Coptis japonica* | Coptidis Rhizoma | Ranunculaceae |
| **M0004375** | *Coptis deltoidea* | Coptidis Rhizoma | Ranunculaceae |
| **M0004376** | *Coptis teetoides* | Coptidis Rhizoma | Ranunculaceae |
| **M0004377** | *Coptis japonica var. dissecta* | Coptidis Rhizoma | Ranunculaceae |
| **M0004379** | *Coptis japonica* | Alcohol cum Coptidis Rhizoma | Ranunculaceae |
| **M0004380** | *Coptis chinensis* | Alcohol cum Coptidis Rhizoma | Ranunculaceae |
| **M0004387** | *Phellodendron chinense* | Phellodendri Cortex | Rutaceae |
| **M0004394** | *Phellodendron amurense* | Phellodendri Radix | Rutaceae |

**Table S5**. List of plants illustrated in **Figures 2G** and **2H**.

| **Plant ID** | **Scientific Name** | **Latin Name** | **Family** |
| --- | --- | --- | --- |
| **M0000112** | *Prunus humilis* | Pruni Humilis Semen | Rosaceae |
| **M0000501** | *Apium graveolens var. dulce* | Apii Herba | Umbelliferae |
| **M0001048** | *Lasiosphaera nipponica* | Lasiosphaera Seu Calvatia | Lycoperdaceae |
| **M0002838** | *Prunus humilis* | Pruni Cortex | Rosaceae |
| **M0002840** | *Prunus humilis* | Pruni Folium | Rosaceae |
| **M0003141** | *Amaranthus mangostanus* | Amaranthi Herba | Amaranthaceae |
| **M0003628** | *Prunus humilis* | Pruni Lignum | Rosaceae |
| **M0003833** | *Allium fistulosum* | Allii Radix | Liliaceae |
| **M0003836** | *Allium fistulosum* | Allii Fistulosi Succus | Liliaceae |
| **M0003852** | *Ailanthus altissima\|Ailanthus glandulosa* | Ailanthi Folium | Simaroubaceae |
| **M0004209** | *Amaranthus mangostanus* | Amaranthi Semen | Amaranthaceae |

**Table S6.** List of chemicals which are targeted on phosphodiesterase 10A in **Figure 3A**.

| **Chemical ID** | **Chemical Name** | **inchikey** | **Chemical Class** | | | | |
| --- | --- | --- | --- | --- | --- | --- | --- |
|  |  |  | **Kingdom** | **Superclass** | **Class** | **Subclass** | **Direct_parent** |
| **1570** | 1-(2'-Hydroxyethyl)-4-methoxy-beta-carboline | XAKXPUZLFGDHAO-UHFFFAOYSA-N | Organic compounds | Alkaloids and derivatives | Harmala alkaloids | None | Harmala alkaloids |
| **5411** | 1-Hydroxycanthin-6-one | LWYFITNQEPSUDK-UHFFFAOYSA-N | Organic compounds | Alkaloids and derivatives | Indolonaphthyridine alkaloids | None | Indolonaphthyridine alkaloids |
| **1574** | 1-Methoxycanthin-6-one | LEPXKGXTXIACRO-UHFFFAOYSA-N | Organic compounds | Alkaloids and derivatives | Indolonaphthyridine alkaloids | None | Indolonaphthyridine alkaloids |
| **17289** | 1-Hydroxycanthin-6-one; Me ether | LEPXKGXTXIACRO-UHFFFAOYSA-N | Organic compounds | Alkaloids and derivatives | Indolonaphthyridine alkaloids | None | Indolonaphthyridine alkaloids |
| **537** | Canthin-6-one | ZERVJPYNQLONEK-UHFFFAOYSA-N | Organic compounds | Alkaloids and derivatives | Indolonaphthyridine alkaloids | None | Indolonaphthyridine alkaloids |
| **629** | 2-Ethyl-3-methylpyrazine | LNIMMWYNSBZESE-UHFFFAOYSA-N | Organic compounds | Organoheterocyclic compounds | Diazines | Pyrazines | Pyrazines |
| **1558** | 2-Hydroxycanthin-6-one | BDCOKQCMZJCYDY-UHFFFAOYSA-N | Organic compounds | Alkaloids and derivatives | Indolonaphthyridine alkaloids | None | Indolonaphthyridine alkaloids |
| **631** | 2-Methylpyrazine | CAWHJQAVHZEVTJ-UHFFFAOYSA-N | Organic compounds | Organoheterocyclic compounds | Diazines | Pyrazines | Pyrazines |
| **632** | 2-Methoxy-3-methylpyrazine | VKJIAEQRKBQLLA-UHFFFAOYSA-N | Organic compounds | Organoheterocyclic compounds | Diazines | Pyrazines | Methoxypyrazines |
| **1561** | 5-Hydroxycanthin-6-one | HOTCTNQHGRFRLK-UHFFFAOYSA-N | Organic compounds | Alkaloids and derivatives | Indolonaphthyridine alkaloids | None | Indolonaphthyridine alkaloids |
| **630** | 2-Ethylpyrazine | KVFIJIWMDBAGDP-UHFFFAOYSA-N | Organic compounds | Organoheterocyclic compounds | Diazines | Pyrazines | Pyrazines |
| **1564** | 5-Hydroxymethylcanthin-6-one | NGLFCPGASUBUBB-UHFFFAOYSA-N | Organic compounds | Alkaloids and derivatives | Indolonaphthyridine alkaloids | None | Indolonaphthyridine alkaloids |

**Table S7.** List of chemicals linked with the plant, *Morus alba* and the target, tyrosinase.

| **Chemical ID** | **Chemical Name** | **inchikey** | **Chemical Class** | | | | |
| --- | --- | --- | --- | --- | --- | --- | --- |
|  |  |  | **Kingdom** | **Superclass** | **Class** | **Subclass** | **Direct_parent** |
| **7808** | Mulberrofuran A | MQYYTNPXQXSQGM-CAOOACKPSA-N | Organic compounds | Phenylpropanoids and polyketides | 2-arylbenzofuran flavonoids | None | 2-arylbenzofuran flavonoids |
| **6657** | Morachalcone A | NXBYIJSAISXPKJ-WEVVVXLNSA-N | Organic compounds | Phenylpropanoids and polyketides | Linear 1,3-diarylpropanoids | Chalcones and dihydrochalcones | 3-prenylated chalcones |
| **7809** | Resorcinol | GHMLBKRAJCXXBS-UHFFFAOYSA-N | Organic compounds | Benzenoids | Phenols | Benzenediols | Resorcinols |
| **7810** | Oxydihydroresveratrol | IEOZKGCYMAJAHS-UHFFFAOYSA-N | Organic compounds | Phenylpropanoids and polyketides | Stilbenes | None | Stilbenes |
| **7811** | Dihydroresveratrol | HITJFUSPLYBJPE-UHFFFAOYSA-N | Organic compounds | Phenylpropanoids and polyketides | Stilbenes | None | Stilbenes |
| **7812** | Oxyresveratrol (E-form) | PDHAOJSHSJQANO-OWOJBTEDSA-N | Organic compounds | Phenylpropanoids and polyketides | Stilbenes | None | Stilbenes |
| **7813** | beta-Resorcylaldehyde | IUNJCFABHJZSKB-UHFFFAOYSA-N | Organic compounds | Organic oxygen compounds | Organooxygen compounds | Carbonyl compounds | Hydroxybenzaldehydes |
| **788** | Albafuran A | KGOOVUKZICPAIZ-FRKPEAEDSA-N | Organic compounds | Phenylpropanoids and polyketides | 2-arylbenzofuran flavonoids | None | 2-arylbenzofuran flavonoids |
| **789** | Albafuran B | ODASNNUHHLRPEL-FRKPEAEDSA-N | Organic compounds | Phenylpropanoids and polyketides | 2-arylbenzofuran flavonoids | None | 2-arylbenzofuran flavonoids |
| **791** | Albanol A | MJJWBJFYYRAYKU-UHFFFAOYSA-N | Organic compounds | Phenylpropanoids and polyketides | 2-arylbenzofuran flavonoids | None | 2-arylbenzofuran flavonoids |
| **792** | 5,7-Dihydroxychromone | NYCXYKOXLNBYID-UHFFFAOYSA-N | Organic compounds | Organoheterocyclic compounds | Benzopyrans | 1-benzopyrans | Chromones |
| **793** | Dimoracin | GKHRLTCUMXVTAV-UHFFFAOYSA-N | Organic compounds | Phenylpropanoids and polyketides | 2-arylbenzofuran flavonoids | None | 2-arylbenzofuran flavonoids |
| **17947** | 1-(2,4-Dihydroxyphenyl)-2-(3,5-dihydroxyphenyl)ethylene; 3'-O-beta-D-Glucopyranoside | GGQVPULXXVQLRT-OWOJBTEDSA-N | Organic compounds | Phenylpropanoids and polyketides | Stilbenes | Stilbene glycosides | Stilbene glycosides |
| **796** | Moracin C | ZTGHWUWBQNCCOH-UHFFFAOYSA-N | Organic compounds | Phenylpropanoids and polyketides | 2-arylbenzofuran flavonoids | None | 2-arylbenzofuran flavonoids |
| **797** | Mulberrofuran F | SCNZCLDHJJSZBK-UHFFFAOYSA-N | Organic compounds | Phenylpropanoids and polyketides | 2-arylbenzofuran flavonoids | None | 2-arylbenzofuran flavonoids |
| **17948** | cis-Mulberroside A | HPSWAEGGWLOOKT-UPHRSURJSA-N | Organic compounds | Phenylpropanoids and polyketides | Stilbenes | Stilbene glycosides | Stilbene glycosides |
| **799** | Mulberrofuran P | WZXFDNVWWQJWRC-UHFFFAOYSA-N | Organic compounds | Phenylpropanoids and polyketides | 2-arylbenzofuran flavonoids | None | 2-arylbenzofuran flavonoids |
| **7337** | Mulberranol | FMKONXHUEWRDEL-UHFFFAOYSA-N | Organic compounds | Phenylpropanoids and polyketides | Flavonoids | Flavones | 6-prenylated flavones |
| **8623** | Resveratrol | LUKBXSAWLPMMSZ-OWOJBTEDSA-N | Organic compounds | Phenylpropanoids and polyketides | Stilbenes | None | Stilbenes |
| **8369** | Broussonin A | MSNVBURPCQDLEP-UHFFFAOYSA-N | Organic compounds | Phenylpropanoids and polyketides | Linear 1,3-diarylpropanoids | Cinnamylphenols | Cinnamylphenols |
| **8370** | Broussonin B | CJJJQWAYMRTLJT-UHFFFAOYSA-N | Organic compounds | Phenylpropanoids and polyketides | Linear 1,3-diarylpropanoids | Cinnamylphenols | Cinnamylphenols |
| **823** | Moracin M | LHPRYOJTASOZGJ-UHFFFAOYSA-N | Organic compounds | Phenylpropanoids and polyketides | 2-arylbenzofuran flavonoids | None | 2-arylbenzofuran flavonoids |
| **8887** | Kuwanon Y | YYUHPJKWIHNMSV-NSCUHMNNSA-N | Organic compounds | Phenylpropanoids and polyketides | Diarylheptanoids | Linear diarylheptanoids | Linear diarylheptanoids |
| **1215** | omega-Hydroxymoracin N | GPKLNIVEGYWQJZ-BIIKFXOESA-N | Organic compounds | Phenylpropanoids and polyketides | 2-arylbenzofuran flavonoids | None | 2-arylbenzofuran flavonoids |
| **1225** | Moracin N | WBSCSIABHGPAMC-UHFFFAOYSA-N | Organic compounds | Phenylpropanoids and polyketides | 2-arylbenzofuran flavonoids | None | 2-arylbenzofuran flavonoids |
| **10575** | Dihydrooxyresveratrol | IEOZKGCYMAJAHS-UHFFFAOYSA-N | Organic compounds | Phenylpropanoids and polyketides | Stilbenes | None | Stilbenes |
| **6617** | Mulberrin | UWQYBLOHTQWSQD-UHFFFAOYSA-N | Organic compounds | Phenylpropanoids and polyketides | Flavonoids | Flavones | 8-prenylated flavones |
| **6619** | Kuwanon A | DBUNRZUFILGKHP-UHFFFAOYSA-N | Organic compounds | Phenylpropanoids and polyketides | Flavonoids | Flavones | 3-prenylated flavones |
| **6238** | Chrysanthemin | RKWHWFONKJEUEF-UHFFFAOYSA-O | Organic compounds | Phenylpropanoids and polyketides | Flavonoids | Flavonoid glycosides | Anthocyanidin-3-O-glycosides |
| **6624** | Kuwanon G | APPXYONGBIXGRO-UHFFFAOYSA-N | Organic compounds | Phenylpropanoids and polyketides | Flavonoids | Flavones | 8-prenylated flavones |
| **6625** | Kuwanon H | DKBPTKFKCCNXNH-UHFFFAOYSA-N | Organic compounds | Phenylpropanoids and polyketides | Flavonoids | Flavones | 8-prenylated flavones |
| **6626** | Kuwanon I | KBAPHKOHTBBCTO-AWNIVKPZSA-N | Organic compounds | Phenylpropanoids and polyketides | Diarylheptanoids | Linear diarylheptanoids | Linear diarylheptanoids |
| **6627** | Kuwanon J | KBAPHKOHTBBCTO-AWNIVKPZSA-N | Organic compounds | Phenylpropanoids and polyketides | Diarylheptanoids | Linear diarylheptanoids | Linear diarylheptanoids |
| **6628** | Kuwanon R | LSWPUMCBBKEXMW-VIZOYTHASA-N | Organic compounds | Phenylpropanoids and polyketides | Diarylheptanoids | Linear diarylheptanoids | Linear diarylheptanoids |
| **6629** | Kuwanon Q | WDCSNUYKXLXPBM-OVCLIPMQSA-N | Organic compounds | Phenylpropanoids and polyketides | Diarylheptanoids | Linear diarylheptanoids | Linear diarylheptanoids |
| **6630** | Kuwanon V | SUQUIVSLHDOSQP-REZTVBANSA-N | Organic compounds | Phenylpropanoids and polyketides | Diarylheptanoids | Linear diarylheptanoids | Linear diarylheptanoids |
| **6631** | Kuwanon K | FEAIYAOTVUYWQZ-UHFFFAOYSA-N | Organic compounds | Phenylpropanoids and polyketides | Flavonoids | Flavones | 3'-prenylated flavones |
| **6633** | Albafuran C | SEUPIEHHWMMMQG-UHFFFAOYSA-N | Organic compounds | Phenylpropanoids and polyketides | 2-arylbenzofuran flavonoids | None | 2-arylbenzofuran flavonoids |
| **6636** | Morusinol | AFOKZNPZDXHDHD-UHFFFAOYSA-N | Organic compounds | Phenylpropanoids and polyketides | Flavonoids | Flavones | 3-prenylated flavones |
| **6637** | Mulberrofuran E | RQIKMRKKKIMUNB-UHFFFAOYSA-N | Organic compounds | Phenylpropanoids and polyketides | 2-arylbenzofuran flavonoids | None | 2-arylbenzofuran flavonoids |
| **6638** | Mulberrofuran T | XMXZFZDVDCIFKB-UHFFFAOYSA-N | Organic compounds | Phenylpropanoids and polyketides | 2-arylbenzofuran flavonoids | None | 2-arylbenzofuran flavonoids |
| **6639** | Mulberrofuran C | WTGKDESIYCVAOP-UHFFFAOYSA-N | Organic compounds | Phenylpropanoids and polyketides | 2-arylbenzofuran flavonoids | None | 2-arylbenzofuran flavonoids |
| **6641** | Sanggenon E | NJQZTHTXKPCAHD-UHFFFAOYSA-N | Organic compounds | Phenylpropanoids and polyketides | Stilbenes | None | Stilbenes |
| **6643** | Sanggenon P | CUJJTBMGUHNKPO-UHFFFAOYSA-N | Organic compounds | Phenylpropanoids and polyketides | Diarylheptanoids | Linear diarylheptanoids | Linear diarylheptanoids |
| **6644** | Chalcomoracin | SEHVRKPXIDOTRX-UHFFFAOYSA-N | Organic compounds | Phenylpropanoids and polyketides | 2-arylbenzofuran flavonoids | None | 2-arylbenzofuran flavonoids |
| **6646** | Morusin | XFFOMNJIDRDDLQ-UHFFFAOYSA-N | Organic compounds | Phenylpropanoids and polyketides | Flavonoids | Flavones | 3-prenylated flavones |
| **6647** | Cyclomulberrin | SYFDWXWLRGHYAJ-UHFFFAOYSA-N | Organic compounds | Phenylpropanoids and polyketides | Flavonoids | Pyranoflavonoids | Pyranoflavonoids |
| **6648** | Moracenin D | PYXAHGLTSLDDJH-UHFFFAOYSA-N | Organic compounds | Phenylpropanoids and polyketides | Flavonoids | Flavones | 8-prenylated flavones |
| **7806** | 2,4-Dihydroxybenzoic acid ethyl ether | BRDIPNLKURUXCU-UHFFFAOYSA-N | Organic compounds | Benzenoids | Benzene and substituted derivatives | Benzoic acids and derivatives | p-Hydroxybenzoic acid alkyl esters |
| **7807** | Kuwanol E | HGFWVFTZYRJFRI-AATRIKPKSA-N | Organic compounds | Phenylpropanoids and polyketides | Diarylheptanoids | Linear diarylheptanoids | Linear diarylheptanoids |

**Table S8.** List of chemicals linked with the plants, *Carthamus tinctorius*, *Cudrania tricuspidata*, *Glycyrrhiza inflata* and *Glycyrrhiza uralensis* and the target, tyrosinase.

| **Chemical ID** | **Chemical Name** | **inchikey** | **Chemical Class** | | | | |
| --- | --- | --- | --- | --- | --- | --- | --- |
|  |  |  | **Kingdom** | **Superclass** | **Class** | **Subclass** | **Direct_parent** |
| ***Carthamus tinctorius*** | | | | | | | |
| **7232** | Safflomin C | CCEKPTFNQKNHKZ-XCVCLJGOSA-N | Organic compounds | Phenylpropanoids and polyketides | Diarylheptanoids | Linear diarylheptanoids | Curcuminoids |
| **6187** | Safflor Yellow B | WBKUQZJRZIYAJL-YDWXAUTNSA-N | None | None | None | None | None |
| **17516** | PRE | ITBPOPSLMGJTQH-YDWXAUTNSA-N | None | None | None | None | None |
| **7231** | Safflomin A | NNXHCBKOBDDJFM-ZZXKWVIFSA-N | None | None | None | None | None |
| **920** | N-(p-Coumaroyl)-5-hydroxytryptamine | WLZPAFGVOWCVMG-FPYGCLRLSA-N | Organic compounds | Organoheterocyclic compounds | Indoles and derivatives | Tryptamines and derivatives | N-acylserotonins |
| **918** | N-Feruloylserotonin | WGHKJYWENWLOMY-XVNBXDOJSA-N | Organic compounds | Organoheterocyclic compounds | Indoles and derivatives | Tryptamines and derivatives | N-acylserotonins |
| **17495** | 4,4''-Bi(N-4-hydroxycinnamoylserotonin); (E,E)-form | OIUNULHHOJRJBI-IAGONARPSA-N | Organic compounds | Organoheterocyclic compounds | Indoles and derivatives | Tryptamines and derivatives | N-acylserotonins |
| **17496** | 4-[N-(p-Coumaroyl)serotonin-4''-yl]-N-feruloylserotonin | ZPNFTINOYMQICL-PWSZKDBUSA-N | Organic compounds | Organoheterocyclic compounds | Indoles and derivatives | Tryptamines and derivatives | N-acylserotonins |
| **17497** | 4,4''-Bis(N-feruloyl)serotonin | URHRFQQPWDNDQE-ACFHMISVSA-N | Organic compounds | Organoheterocyclic compounds | Indoles and derivatives | Tryptamines and derivatives | N-acylserotonins |
| **7261** | Safflor yellow A | YKDBQWHZEDSTAI-ZZXKWVIFSA-N | Organic compounds | Phenylpropanoids and polyketides | Cinnamic acids and derivatives | Hydroxycinnamic acids and derivatives | Hydroxycinnamic acids and derivatives |
| **1502** | N-(p-Coumaroyl)serotonin | WLZPAFGVOWCVMG-FPYGCLRLSA-N | Organic compounds | Organoheterocyclic compounds | Indoles and derivatives | Tryptamines and derivatives | N-acylserotonins |
| **1503** | N-(p-Coumaroyl)serotonin mono-bera-D-glucopyranoside | LPGWQGDUKIPAME-FPYGCLRLSA-N | Organic compounds | Organic oxygen compounds | Organooxygen compounds | Carbohydrates and carbohydrate conjugates | Phenolic glycosides |
| ***Cudrania tricuspidata*** | | | | | | | |
| **7808** | Mulberrofuran A | MQYYTNPXQXSQGM-CAOOACKPSA-N | Organic compounds | Phenylpropanoids and polyketides | 2-arylbenzofuran flavonoids | None | 2-arylbenzofuran flavonoids |
| **6657** | Morachalcone A | NXBYIJSAISXPKJ-WEVVVXLNSA-N | Organic compounds | Phenylpropanoids and polyketides | Linear 1,3-diarylpropanoids | Chalcones and dihydrochalcones | 3-prenylated chalcones |
| **7810** | Oxydihydroresveratrol | IEOZKGCYMAJAHS-UHFFFAOYSA-N | Organic compounds | Phenylpropanoids and polyketides | Stilbenes | None | Stilbenes |
| **7811** | Dihydroresveratrol | HITJFUSPLYBJPE-UHFFFAOYSA-N | Organic compounds | Phenylpropanoids and polyketides | Stilbenes | None | Stilbenes |
| **7812** | Oxyresveratrol (E-form) | PDHAOJSHSJQANO-OWOJBTEDSA-N | Organic compounds | Phenylpropanoids and polyketides | Stilbenes | None | Stilbenes |
| **7813** | beta-Resorcylaldehyde | IUNJCFABHJZSKB-UHFFFAOYSA-N | Organic compounds | Organic oxygen compounds | Organooxygen compounds | Carbonyl compounds | Hydroxybenzaldehydes |
| **7809** | Resorcinol | GHMLBKRAJCXXBS-UHFFFAOYSA-N | Organic compounds | Benzenoids | Phenols | Benzenediols | Resorcinols |
| **788** | Albafuran A | KGOOVUKZICPAIZ-FRKPEAEDSA-N | Organic compounds | Phenylpropanoids and polyketides | 2-arylbenzofuran flavonoids | None | 2-arylbenzofuran flavonoids |
| **789** | Albafuran B | ODASNNUHHLRPEL-FRKPEAEDSA-N | Organic compounds | Phenylpropanoids and polyketides | 2-arylbenzofuran flavonoids | None | 2-arylbenzofuran flavonoids |
| **791** | Albanol A | MJJWBJFYYRAYKU-UHFFFAOYSA-N | Organic compounds | Phenylpropanoids and polyketides | 2-arylbenzofuran flavonoids | None | 2-arylbenzofuran flavonoids |
| **792** | 5,7-Dihydroxychromone | NYCXYKOXLNBYID-UHFFFAOYSA-N | Organic compounds | Organoheterocyclic compounds | Benzopyrans | 1-benzopyrans | Chromones |
| **793** | Dimoracin | GKHRLTCUMXVTAV-UHFFFAOYSA-N | Organic compounds | Phenylpropanoids and polyketides | 2-arylbenzofuran flavonoids | None | 2-arylbenzofuran flavonoids |
| **17947** | 1-(2,4-Dihydroxyphenyl)-2-(3,5-dihydroxyphenyl)ethylene; 3'-O-beta-D-Glucopyranoside | GGQVPULXXVQLRT-OWOJBTEDSA-N | Organic compounds | Phenylpropanoids and polyketides | Stilbenes | Stilbene glycosides | Stilbene glycosides |
| **17948** | cis-Mulberroside A | HPSWAEGGWLOOKT-UPHRSURJSA-N | Organic compounds | Phenylpropanoids and polyketides | Stilbenes | Stilbene glycosides | Stilbene glycosides |
| **796** | Moracin C | ZTGHWUWBQNCCOH-UHFFFAOYSA-N | Organic compounds | Phenylpropanoids and polyketides | 2-arylbenzofuran flavonoids | None | 2-arylbenzofuran flavonoids |
| **797** | Mulberrofuran F | SCNZCLDHJJSZBK-UHFFFAOYSA-N | Organic compounds | Phenylpropanoids and polyketides | 2-arylbenzofuran flavonoids | None | 2-arylbenzofuran flavonoids |
| **799** | Mulberrofuran P | WZXFDNVWWQJWRC-UHFFFAOYSA-N | Organic compounds | Phenylpropanoids and polyketides | 2-arylbenzofuran flavonoids | None | 2-arylbenzofuran flavonoids |
| **7337** | Mulberranol | FMKONXHUEWRDEL-UHFFFAOYSA-N | Organic compounds | Phenylpropanoids and polyketides | Flavonoids | Flavones | 6-prenylated flavones |
| **8623** | Resveratrol | LUKBXSAWLPMMSZ-OWOJBTEDSA-N | Organic compounds | Phenylpropanoids and polyketides | Stilbenes | None | Stilbenes |
| **8369** | Broussonin A | MSNVBURPCQDLEP-UHFFFAOYSA-N | Organic compounds | Phenylpropanoids and polyketides | Linear 1,3-diarylpropanoids | Cinnamylphenols | Cinnamylphenols |
| **8370** | Broussonin B | CJJJQWAYMRTLJT-UHFFFAOYSA-N | Organic compounds | Phenylpropanoids and polyketides | Linear 1,3-diarylpropanoids | Cinnamylphenols | Cinnamylphenols |
| **823** | Moracin M | LHPRYOJTASOZGJ-UHFFFAOYSA-N | Organic compounds | Phenylpropanoids and polyketides | 2-arylbenzofuran flavonoids | None | 2-arylbenzofuran flavonoids |
| **8887** | Kuwanon Y | YYUHPJKWIHNMSV-NSCUHMNNSA-N | Organic compounds | Phenylpropanoids and polyketides | Diarylheptanoids | Linear diarylheptanoids | Linear diarylheptanoids |
| **1215** | omega-Hydroxymoracin N | GPKLNIVEGYWQJZ-BIIKFXOESA-N | Organic compounds | Phenylpropanoids and polyketides | 2-arylbenzofuran flavonoids | None | 2-arylbenzofuran flavonoids |
| **1225** | Moracin N | WBSCSIABHGPAMC-UHFFFAOYSA-N | Organic compounds | Phenylpropanoids and polyketides | 2-arylbenzofuran flavonoids | None | 2-arylbenzofuran flavonoids |
| **10575** | Dihydrooxyresveratrol | IEOZKGCYMAJAHS-UHFFFAOYSA-N | Organic compounds | Phenylpropanoids and polyketides | Stilbenes | None | Stilbenes |
| **6617** | Mulberrin | UWQYBLOHTQWSQD-UHFFFAOYSA-N | Organic compounds | Phenylpropanoids and polyketides | Flavonoids | Flavones | 8-prenylated flavones |
| **6619** | Kuwanon A | DBUNRZUFILGKHP-UHFFFAOYSA-N | Organic compounds | Phenylpropanoids and polyketides | Flavonoids | Flavones | 3-prenylated flavones |
| **6238** | Chrysanthemin | RKWHWFONKJEUEF-UHFFFAOYSA-O | Organic compounds | Phenylpropanoids and polyketides | Flavonoids | Flavonoid glycosides | Anthocyanidin-3-O-glycosides |
| **6624** | Kuwanon G | APPXYONGBIXGRO-UHFFFAOYSA-N | Organic compounds | Phenylpropanoids and polyketides | Flavonoids | Flavones | 8-prenylated flavones |
| **6625** | Kuwanon H | DKBPTKFKCCNXNH-UHFFFAOYSA-N | Organic compounds | Phenylpropanoids and polyketides | Flavonoids | Flavones | 8-prenylated flavones |
| **6626** | Kuwanon I | KBAPHKOHTBBCTO-AWNIVKPZSA-N | Organic compounds | Phenylpropanoids and polyketides | Diarylheptanoids | Linear diarylheptanoids | Linear diarylheptanoids |
| **6627** | Kuwanon J | KBAPHKOHTBBCTO-AWNIVKPZSA-N | Organic compounds | Phenylpropanoids and polyketides | Diarylheptanoids | Linear diarylheptanoids | Linear diarylheptanoids |
| **6628** | Kuwanon R | LSWPUMCBBKEXMW-VIZOYTHASA-N | Organic compounds | Phenylpropanoids and polyketides | Diarylheptanoids | Linear diarylheptanoids | Linear diarylheptanoids |
| **6629** | Kuwanon Q | WDCSNUYKXLXPBM-OVCLIPMQSA-N | Organic compounds | Phenylpropanoids and polyketides | Diarylheptanoids | Linear diarylheptanoids | Linear diarylheptanoids |
| **6630** | Kuwanon V | SUQUIVSLHDOSQP-REZTVBANSA-N | Organic compounds | Phenylpropanoids and polyketides | Diarylheptanoids | Linear diarylheptanoids | Linear diarylheptanoids |
| **6631** | Kuwanon K | FEAIYAOTVUYWQZ-UHFFFAOYSA-N | Organic compounds | Phenylpropanoids and polyketides | Flavonoids | Flavones | 3'-prenylated flavones |
| **6633** | Albafuran C | SEUPIEHHWMMMQG-UHFFFAOYSA-N | Organic compounds | Phenylpropanoids and polyketides | 2-arylbenzofuran flavonoids | None | 2-arylbenzofuran flavonoids |
| **6636** | Morusinol | AFOKZNPZDXHDHD-UHFFFAOYSA-N | Organic compounds | Phenylpropanoids and polyketides | Flavonoids | Flavones | 3-prenylated flavones |
| **6637** | Mulberrofuran E | RQIKMRKKKIMUNB-UHFFFAOYSA-N | Organic compounds | Phenylpropanoids and polyketides | 2-arylbenzofuran flavonoids | None | 2-arylbenzofuran flavonoids |
| **6638** | Mulberrofuran T | XMXZFZDVDCIFKB-UHFFFAOYSA-N | Organic compounds | Phenylpropanoids and polyketides | 2-arylbenzofuran flavonoids | None | 2-arylbenzofuran flavonoids |
| **6639** | Mulberrofuran C | WTGKDESIYCVAOP-UHFFFAOYSA-N | Organic compounds | Phenylpropanoids and polyketides | 2-arylbenzofuran flavonoids | None | 2-arylbenzofuran flavonoids |
| **6641** | Sanggenon E | NJQZTHTXKPCAHD-UHFFFAOYSA-N | Organic compounds | Phenylpropanoids and polyketides | Stilbenes | None | Stilbenes |
| **6643** | Sanggenon P | CUJJTBMGUHNKPO-UHFFFAOYSA-N | Organic compounds | Phenylpropanoids and polyketides | Diarylheptanoids | Linear diarylheptanoids | Linear diarylheptanoids |
| **6644** | Chalcomoracin | SEHVRKPXIDOTRX-UHFFFAOYSA-N | Organic compounds | Phenylpropanoids and polyketides | 2-arylbenzofuran flavonoids | None | 2-arylbenzofuran flavonoids |
| **6646** | Morusin | XFFOMNJIDRDDLQ-UHFFFAOYSA-N | Organic compounds | Phenylpropanoids and polyketides | Flavonoids | Flavones | 3-prenylated flavones |
| **6647** | Cyclomulberrin | SYFDWXWLRGHYAJ-UHFFFAOYSA-N | Organic compounds | Phenylpropanoids and polyketides | Flavonoids | Pyranoflavonoids | Pyranoflavonoids |
| **6648** | Moracenin D | PYXAHGLTSLDDJH-UHFFFAOYSA-N | Organic compounds | Phenylpropanoids and polyketides | Flavonoids | Flavones | 8-prenylated flavones |
| **7806** | 2,4-Dihydroxybenzoic acid ethyl ether | BRDIPNLKURUXCU-UHFFFAOYSA-N | Organic compounds | Benzenoids | Benzene and substituted derivatives | Benzoic acids and derivatives | p-Hydroxybenzoic acid alkyl esters |
| **7807** | Kuwanol E | HGFWVFTZYRJFRI-AATRIKPKSA-N | Organic compounds | Phenylpropanoids and polyketides | Diarylheptanoids | Linear diarylheptanoids | Linear diarylheptanoids |
| ***Glycyrrhiza inflata*** | | | | | | | |
| **17792** | Licochalcone C | WBDNTJSRHDSPSR-KPKJPENVSA-N | Organic compounds | Phenylpropanoids and polyketides | Linear 1,3-diarylpropanoids | Chalcones and dihydrochalcones | Retrochalcones |
| **7300** | 5'-Prenyllicodione | VAWLLIOUAFRMHN-UHFFFAOYSA-N | Organic compounds | Phenylpropanoids and polyketides | Linear 1,3-diarylpropanoids | Chalcones and dihydrochalcones | Retro-dihydrochalcones |
| **7301** | Glycyrdione A | VDOHBGQSFOWYTB-UHFFFAOYSA-N | Organic compounds | Phenylpropanoids and polyketides | Linear 1,3-diarylpropanoids | Chalcones and dihydrochalcones | Retro-dihydrochalcones |
| **7303** | Glyinflanin A | VDOHBGQSFOWYTB-UHFFFAOYSA-N | Organic compounds | Phenylpropanoids and polyketides | Linear 1,3-diarylpropanoids | Chalcones and dihydrochalcones | Retro-dihydrochalcones |
| **7317** | Licochalcone C | WBDNTJSRHDSPSR-KPKJPENVSA-N | Organic compounds | Phenylpropanoids and polyketides | Linear 1,3-diarylpropanoids | Chalcones and dihydrochalcones | Retrochalcones |
| **7318** | Licochalcone D | RETRVWFVEFCGOK-RMKNXTFCSA-N | Organic compounds | Phenylpropanoids and polyketides | Linear 1,3-diarylpropanoids | Chalcones and dihydrochalcones | 3-prenylated chalcones |
| **5960** | Isoliquiritigenin | DXDRHHKMWQZJHT-FPYGCLRLSA-N | Organic compounds | Phenylpropanoids and polyketides | Linear 1,3-diarylpropanoids | Chalcones and dihydrochalcones | 2'-Hydroxychalcones |
| **8137** | Glycycoumarin | NZYSZZDSYIBYLC-UHFFFAOYSA-N | Organic compounds | Phenylpropanoids and polyketides | Isoflavonoids | Hydroxyisoflavonoids | Hydroxyisoflavonoids |
| **10318** | Licoflavone B | GLDVIKFETPAZNV-UHFFFAOYSA-N | Organic compounds | Phenylpropanoids and polyketides | Flavonoids | Flavones | 6-prenylated flavones |
| **10330** | Licoflavone C | MEHHCBRCXIDGKZ-UHFFFAOYSA-N | Organic compounds | Phenylpropanoids and polyketides | Flavonoids | Flavones | 8-prenylated flavones |
| **5980** | Echinatin | QJKMIJNRNRLQSS-WEVVVXLNSA-N | Organic compounds | Phenylpropanoids and polyketides | Linear 1,3-diarylpropanoids | Chalcones and dihydrochalcones | Retrochalcones |
| **5981** | Licochalcone A | KAZSKMJFUPEHHW-DHZHZOJOSA-N | Organic compounds | Phenylpropanoids and polyketides | Linear 1,3-diarylpropanoids | Chalcones and dihydrochalcones | Retrochalcones |
| **5982** | Licochalcone B | DRDRYGIIYOPBBZ-XBXARRHUSA-N | Organic compounds | Phenylpropanoids and polyketides | Linear 1,3-diarylpropanoids | Chalcones and dihydrochalcones | Retrochalcones |
| **5983** | Isoliquiritin | YNWXJFQOCHMPCK-FPYGCLRLSA-N | Organic compounds | Phenylpropanoids and polyketides | Flavonoids | Flavonoid glycosides | Flavonoid O-glycosides |
| **17774** | Licoflavone B | GLDVIKFETPAZNV-UHFFFAOYSA-N | Organic compounds | Phenylpropanoids and polyketides | Flavonoids | Flavones | 6-prenylated flavones |
| **17777** | Glyinflanin I; (R)-form | AFQCFVZKNRARLS-UHFFFAOYSA-N | Organic compounds | Phenylpropanoids and polyketides | Isoflavonoids | Pyranoisoflavonoids | Pyranoisoflavonoids |
| **17781** | Glyinflanin E | YSMQGBBJPVUXEX-UHFFFAOYSA-N | Organic compounds | Phenylpropanoids and polyketides | Linear 1,3-diarylpropanoids | Chalcones and dihydrochalcones | Retro-dihydrochalcones |
| **7162** | Licoflavone A | HJGURBGBPIKRER-UHFFFAOYSA-N | Organic compounds | Phenylpropanoids and polyketides | Flavonoids | Flavones | 6-prenylated flavones |
| **17787** | Inflacoumarin A | RNBLSJGPSGNSIN-UHFFFAOYSA-N | Organic compounds | Phenylpropanoids and polyketides | Neoflavonoids | Prenylated neoflavonoids | Prenylated neoflavonoids |
| **17791** | Licochalcone D | RETRVWFVEFCGOK-RMKNXTFCSA-N | Organic compounds | Phenylpropanoids and polyketides | Linear 1,3-diarylpropanoids | Chalcones and dihydrochalcones | 3-prenylated chalcones |
| ***Glycyrrhiza uralensis*** | | | | | | | |
| **17793** | Uralenneoside | VWQASRWQZBVNEI-UHFFFAOYSA-N | Organic compounds | Benzenoids | Benzene and substituted derivatives | Benzoic acids and derivatives | p-Hydroxybenzoic acid alkyl esters |
| **17794** | 6''-Acetylliquiritin | HKUBLIRXXFRGKE-UHFFFAOYSA-N | Organic compounds | Phenylpropanoids and polyketides | Flavonoids | Flavonoid glycosides | Flavonoid O-glycosides |
| **17797** | Kanzonol H | JRVDUBFSQWHYRJ-UHFFFAOYSA-N | Organic compounds | Phenylpropanoids and polyketides | Isoflavonoids | O-methylated isoflavonoids | 5-O-methylated isoflavonoids |
| **17799** | Kanzonol L | CLXMHBYPZWNJQI-UHFFFAOYSA-N | Organic compounds | Phenylpropanoids and polyketides | Isoflavonoids | Isoflavans | 6-prenylated isoflavanones |
| **6280** | 2',4,4'-Trihydroxy-3'-prenylchalcone | DUWPGRAKHMEPCM-IZZDOVSWSA-N | Organic compounds | Phenylpropanoids and polyketides | Linear 1,3-diarylpropanoids | Chalcones and dihydrochalcones | 3-prenylated chalcones |
| **17800** | Kanzonol N | ANRYVYXUTFJPOR-UHFFFAOYSA-N | Organic compounds | Phenylpropanoids and polyketides | Isoflavonoids | O-methylated isoflavonoids | 5-O-methylated isoflavonoids |
| **17801** | Kanzonol M | KZQAGRHRFKIERV-UHFFFAOYSA-N | Organic compounds | Phenylpropanoids and polyketides | Isoflavonoids | O-methylated isoflavonoids | 5-O-methylated isoflavonoids |
| **17807** | Kanzonol G | OWVFKLIUBOKWCT-UHFFFAOYSA-N | Organic compounds | Phenylpropanoids and polyketides | Isoflavonoids | Isoflavans | 3'-prenylated isoflavanones |
| **17808** | Kanzonol K | UWUOGPWSIVRQNM-UHFFFAOYSA-N | Organic compounds | Phenylpropanoids and polyketides | Isoflavonoids | Isoflavans | 6-prenylated isoflavanones |
| **17809** | Glyurallin B | DPLWUTYEBRKBLI-UHFFFAOYSA-N | Organic compounds | Phenylpropanoids and polyketides | Isoflavonoids | Isoflav-2-enes | Isoflavones |
| **17810** | Kanzonol P | ZCGOJWAIXQAUMW-UHFFFAOYSA-N | Organic compounds | Phenylpropanoids and polyketides | Isoflavonoids | Furanoisoflavonoids | Pterocarpans |
| **7454** | Gancaonin R | QFAPONVNJTUMHF-UHFFFAOYSA-N | Organic compounds | Phenylpropanoids and polyketides | Stilbenes | None | Stilbenes |
| **7455** | Gancaonin S | VLJOWGMLRJTNDQ-UHFFFAOYSA-N | Organic compounds | Phenylpropanoids and polyketides | Stilbenes | None | Stilbenes |
| **7456** | Gancaonin U | YJJXCOSDPIJFJR-UHFFFAOYSA-N | Organic compounds | Benzenoids | Phenanthrenes and derivatives | Hydrophenanthrenes | Hydrophenanthrenes |
| **7457** | Gancaonin V | UEXOPXIMQJMWKA-UHFFFAOYSA-N | Organic compounds | Benzenoids | Phenanthrenes and derivatives | Hydrophenanthrenes | Hydrophenanthrenes |
| **7205** | Gancaonin C | MEADLGUPYQNUNF-BIIKFXOESA-N | Organic compounds | Phenylpropanoids and polyketides | Isoflavonoids | Isoflav-2-enes | Isoflavones |
| **7206** | Gancaonin D | UCKSAYIMWMIZQJ-QDEBKDIKSA-N | Organic compounds | Phenylpropanoids and polyketides | Isoflavonoids | O-methylated isoflavonoids | 3'-hydroxy,4'-methoxyisoflavonoids |
| **7208** | Gancaonin L | WSOHPJFMARQRFD-UHFFFAOYSA-N | Organic compounds | Phenylpropanoids and polyketides | Isoflavonoids | Isoflav-2-enes | Isoflavones |
| **7209** | Gancaonin M | DDLPIQXHEKZHQX-UHFFFAOYSA-N | Organic compounds | Phenylpropanoids and polyketides | Isoflavonoids | O-methylated isoflavonoids | 4'-O-methylisoflavones |
| **7210** | Gancaonin O | AFJYQKPCJLMHCC-UHFFFAOYSA-N | Organic compounds | Phenylpropanoids and polyketides | Flavonoids | Flavones | 6-prenylated flavones |
| **7212** | Gancaonin Q | WGNIVAMNAWBYRO-UHFFFAOYSA-N | Organic compounds | Phenylpropanoids and polyketides | Flavonoids | Flavones | 6-prenylated flavones |
| **8896** | Gancaonin T | OWBYJSNJVWQEQX-UHFFFAOYSA-N | Organic compounds | Phenylpropanoids and polyketides | Stilbenes | None | Stilbenes |
| **8134** | Glycyrin | FWWGXZYUURXJLK-UHFFFAOYSA-N | Organic compounds | Phenylpropanoids and polyketides | Isoflavonoids | Hydroxyisoflavonoids | Hydroxyisoflavonoids |
| **5960** | Isoliquiritigenin | DXDRHHKMWQZJHT-FPYGCLRLSA-N | Organic compounds | Phenylpropanoids and polyketides | Linear 1,3-diarylpropanoids | Chalcones and dihydrochalcones | 2'-Hydroxychalcones |
| **8137** | Glycycoumarin | NZYSZZDSYIBYLC-UHFFFAOYSA-N | Organic compounds | Phenylpropanoids and polyketides | Isoflavonoids | Hydroxyisoflavonoids | Hydroxyisoflavonoids |
| **5980** | Echinatin | QJKMIJNRNRLQSS-WEVVVXLNSA-N | Organic compounds | Phenylpropanoids and polyketides | Linear 1,3-diarylpropanoids | Chalcones and dihydrochalcones | Retrochalcones |
| **5981** | Licochalcone A | KAZSKMJFUPEHHW-DHZHZOJOSA-N | Organic compounds | Phenylpropanoids and polyketides | Linear 1,3-diarylpropanoids | Chalcones and dihydrochalcones | Retrochalcones |
| **5982** | Licochalcone B | DRDRYGIIYOPBBZ-XBXARRHUSA-N | Organic compounds | Phenylpropanoids and polyketides | Linear 1,3-diarylpropanoids | Chalcones and dihydrochalcones | Retrochalcones |
| **5983** | Isoliquiritin | YNWXJFQOCHMPCK-FPYGCLRLSA-N | Organic compounds | Phenylpropanoids and polyketides | Flavonoids | Flavonoid glycosides | Flavonoid O-glycosides |
| **8542** | 1-O-Protocatechuyl-beta-D-xylose | VWQASRWQZBVNEI-UHFFFAOYSA-N | Organic compounds | Benzenoids | Benzene and substituted derivatives | Benzoic acids and derivatives | p-Hydroxybenzoic acid alkyl esters |
| **5989** | Licoflavonol | TVMHBSODLWMMMV-UHFFFAOYSA-N | Organic compounds | Phenylpropanoids and polyketides | Flavonoids | Flavones | 6-prenylated flavones |
| **5994** | Licoricone | GGWMNTNDTRKETA-UHFFFAOYSA-N | Organic compounds | Phenylpropanoids and polyketides | Isoflavonoids | O-methylated isoflavonoids | 4'-O-methylisoflavones |
| **5996** | Gancaonin A | JQNSUDIGIIGIOL-UHFFFAOYSA-N | Organic compounds | Phenylpropanoids and polyketides | Isoflavonoids | Isoflavans | 6-prenylated isoflavanones |
| **5998** | Gancaonin N | STFVTZQCNYBLNE-UHFFFAOYSA-N | Organic compounds | Phenylpropanoids and polyketides | Isoflavonoids | Isoflavans | 6-prenylated isoflavanones |
| **7162** | Licoflavone A | HJGURBGBPIKRER-UHFFFAOYSA-N | Organic compounds | Phenylpropanoids and polyketides | Flavonoids | Flavones | 6-prenylated flavones |
| **7163** | Lupiwighteone | YGCCASGFIOIXIN-UHFFFAOYSA-N | Organic compounds | Phenylpropanoids and polyketides | Isoflavonoids | Isoflav-2-enes | Isoflavones |

**Table S9.** List of chemicals which are co-targeted on S5R1 and S5R2.

| **Chemical ID** | **Chemical Name** | **inchikey** | **Chemical Class** | | | | |
| --- | --- | --- | --- | --- | --- | --- | --- |
|  |  |  | **Kingdom** | **Superclass** | **Class** | **Subclass** | **Direct_parent** |
| **1925** | Ergost-4-en-3-one | QQIOPZFVTIHASB-UHFFFAOYSA-N | Organic compounds | Lipids and lipid-like molecules | Steroids and steroid derivatives | Ergostane steroids | Ergosterols and derivatives |
| **1926** | Ergost-4-en-3,6-dione | HYALICAWDSDCPS-UHFFFAOYSA-N | Organic compounds | Lipids and lipid-like molecules | Steroids and steroid derivatives | Ergostane steroids | Ergosterols and derivatives |
| **1927** | Stigmast-4-en-3,6-dione | UVFOCYGYACXLAY-UHFFFAOYSA-N | Organic compounds | Lipids and lipid-like molecules | Steroids and steroid derivatives | Stigmastanes and derivatives | Stigmastanes and derivatives |
| **1928** | Stigmasta-4,22-dien-3-one | MKGZDUKUQPPHFM-CMDGGOBGSA-N | Organic compounds | Lipids and lipid-like molecules | Steroids and steroid derivatives | Stigmastanes and derivatives | Stigmastanes and derivatives |
| **10505** | 17-Deoxywithanone | ZTEVDTFJUUJBLP-UHFFFAOYSA-N | Organic compounds | Lipids and lipid-like molecules | Steroids and steroid derivatives | Steroid lactones | Withanolides and derivatives |
| **1929** | Stigmasta-4,22-dien-3,6-dione | XWHBTBBUPBKDBB-CMDGGOBGSA-N | Organic compounds | Lipids and lipid-like molecules | Steroids and steroid derivatives | Stigmastanes and derivatives | Stigmastanes and derivatives |
| **10632** | 24-Ethyl-24-dehydrocholesterol | YYLFLRIUDMIWTD-UHFFFAOYSA-N | Organic compounds | Lipids and lipid-like molecules | Steroids and steroid derivatives | Stigmastanes and derivatives | Stigmastanes and derivatives |
| **10636** | Ergostenone(24R) | QQIOPZFVTIHASB-UHFFFAOYSA-N | Organic compounds | Lipids and lipid-like molecules | Steroids and steroid derivatives | Ergostane steroids | Ergosterols and derivatives |
| **10637** | Stigmasta-4,22-dien-3-one | MKGZDUKUQPPHFM-CMDGGOBGSA-N | Organic compounds | Lipids and lipid-like molecules | Steroids and steroid derivatives | Stigmastanes and derivatives | Stigmastanes and derivatives |
| **10638** | Stigmasta-4,6-dien-3-one | KEAZWUZFBSXOMV-UHFFFAOYSA-N | Organic compounds | Lipids and lipid-like molecules | Steroids and steroid derivatives | Stigmastanes and derivatives | Stigmastanes and derivatives |
| **2702** | Castasterone | VYUIKSFYFRVQLF-UHFFFAOYSA-N | Organic compounds | Lipids and lipid-like molecules | Steroids and steroid derivatives | Bile acids, alcohols and derivatives | Tetrahydroxy bile acids, alcohols and derivatives |
| **9612** | Kusunol | MQWIFDHBNGIVPO-UHFFFAOYSA-N | Organic compounds | Lipids and lipid-like molecules | Prenol lipids | Sesquiterpenoids | Eremophilane, 8,9-secoeremophilane and furoeremophilane sesquiterpenoids |
| **10633** | Stigmasta-5,24(28)-dien-3-ol | OSELKOCHBMDKEJ-QPSGOUHRSA-N | Organic compounds | Lipids and lipid-like molecules | Steroids and steroid derivatives | Stigmastanes and derivatives | Stigmastanes and derivatives |
| **10635** | Cholesta-4,6-dien-3-one | XIWMRKFKSRYSIJ-UHFFFAOYSA-N | Organic compounds | Lipids and lipid-like molecules | Steroids and steroid derivatives | Cholestane steroids | Cholesterols and derivatives |
| **1940** | Isofucosterol | OSELKOCHBMDKEJ-YXSASFKJSA-N | Organic compounds | Lipids and lipid-like molecules | Steroids and steroid derivatives | Stigmastanes and derivatives | Stigmastanes and derivatives |
| **11158** | Dehydroabietinal | YCLCHPWRGSDZKL-UHFFFAOYSA-N | Organic compounds | Lipids and lipid-like molecules | Prenol lipids | Diterpenoids | Diterpenoids |
| **2585** | Panax ginseng glycoside P1 | DAPLORBFRRJHHY-UHFFFAOYSA-N | Organic compounds | Lipids and lipid-like molecules | Steroids and steroid derivatives | Steroidal glycosides | Steroidal glycosides |
| **2330** | alpha-Teresantalic acid | QHDPITNBVDSMQH-UHFFFAOYSA-N | Organic compounds | Lipids and lipid-like molecules | Prenol lipids | Monoterpenoids | Bicyclic monoterpenoids |
| **1956** | 31-Norlanost-8-en-3-ol | IXVNEXDHXGHWLS-UHFFFAOYSA-N | Organic compounds | Lipids and lipid-like molecules | Steroids and steroid derivatives | Cholestane steroids | Cholesterols and derivatives |
| **1959** | 31-Norlanost-9(11)-enol | SZCKXGWHINUNKB-UHFFFAOYSA-N | Organic compounds | Lipids and lipid-like molecules | Steroids and steroid derivatives | Cholestane steroids | Cholesterols and derivatives |
| **2345** | Campesterol | SGNBVLSWZMBQTH-UHFFFAOYSA-N | Organic compounds | Lipids and lipid-like molecules | Steroids and steroid derivatives | Ergostane steroids | Ergosterols and derivatives |
| **1961** | Lophenol | LMYZQUNLYGJIHI-UHFFFAOYSA-N | Organic compounds | Lipids and lipid-like molecules | Steroids and steroid derivatives | Cholestane steroids | Cholesterols and derivatives |
| **3118** | Stigmastane-3,6-dione (5alpha H) | HMMVBUVVQLUGQA-UHFFFAOYSA-N | Organic compounds | Lipids and lipid-like molecules | Steroids and steroid derivatives | Stigmastanes and derivatives | Stigmastanes and derivatives |
| **1967** | 4-alpha-Methylcholest-8-en-3-ol | SCEZIHJVTBQOLS-UHFFFAOYSA-N | Organic compounds | Lipids and lipid-like molecules | Steroids and steroid derivatives | Cholestane steroids | Cholesterols and derivatives |
| **9519** | 5-alpha-Stigmast-22-en-3-one | RTLUSWHIKFIQFU-CMDGGOBGSA-N | Organic compounds | Lipids and lipid-like molecules | Steroids and steroid derivatives | Stigmastanes and derivatives | Stigmastanes and derivatives |
| **9520** | Stigmast-5-en-3-beta-ol-7-one | ICFXJOAKQGDRCT-UHFFFAOYSA-N | Organic compounds | Lipids and lipid-like molecules | Steroids and steroid derivatives | Stigmastanes and derivatives | Stigmastanes and derivatives |
| **1970** | Withanolide B | ZTEVDTFJUUJBLP-UHFFFAOYSA-N | Organic compounds | Lipids and lipid-like molecules | Steroids and steroid derivatives | Steroid lactones | Withanolides and derivatives |
| **2226** | Estradiol | VOXZDWNPVJITMN-UHFFFAOYSA-N | Organic compounds | Lipids and lipid-like molecules | Steroids and steroid derivatives | Estrane steroids | Estrogens and derivatives |
| **9654** | (-)-Ylangene | VLXDPFLIRFYIME-UHFFFAOYSA-N | Organic compounds | Lipids and lipid-like molecules | Prenol lipids | Sesquiterpenoids | Sesquiterpenoids |
| **1847** | (-)-alpha-Copaene | VLXDPFLIRFYIME-UHFFFAOYSA-N | Organic compounds | Lipids and lipid-like molecules | Prenol lipids | Sesquiterpenoids | Sesquiterpenoids |
| **11070** | Aromadendrane-4-beta,10-alpha-diol | DWNPMJOWAWGIMM-UHFFFAOYSA-N | Organic compounds | Lipids and lipid-like molecules | Prenol lipids | Sesquiterpenoids | 5,10-cycloaromadendrane sesquiterpenoids |
| **11076** | Neointermedeol | DPQYOKVMVCQHMY-UHFFFAOYSA-N | Organic compounds | Lipids and lipid-like molecules | Prenol lipids | Sesquiterpenoids | Eudesmane, isoeudesmane or cycloeudesmane sesquiterpenoids |
| **1863** | 3-beta-Hydroxystigmast-5-en-7-one | ICFXJOAKQGDRCT-UHFFFAOYSA-N | Organic compounds | Lipids and lipid-like molecules | Steroids and steroid derivatives | Stigmastanes and derivatives | Stigmastanes and derivatives |
| **2001** | Stigmastan-3-one (5-alpha) | BVVFRHKBULZQCQ-UHFFFAOYSA-N | Organic compounds | Lipids and lipid-like molecules | Steroids and steroid derivatives | Stigmastanes and derivatives | Stigmastanes and derivatives |
| **4562** | 3,5-Acoradiene | ZQMYLSWMANINAL-UHFFFAOYSA-N | Organic compounds | Hydrocarbons | Unsaturated hydrocarbons | Branched unsaturated hydrocarbons | Branched unsaturated hydrocarbons |
| **3032** | Ledol | AYXPYQRXGNDJFU-UHFFFAOYSA-N | Organic compounds | Lipids and lipid-like molecules | Prenol lipids | Sesquiterpenoids | 5,10-cycloaromadendrane sesquiterpenoids |
| **4187** | Stigmasterol | HCXVJBMSMIARIN-CMDGGOBGSA-N | Organic compounds | Lipids and lipid-like molecules | Steroids and steroid derivatives | Stigmastanes and derivatives | Stigmastanes and derivatives |
| **1769** | Stigmast-22-en-3-beta-ol | CSVWWLUMXNHWSU-CMDGGOBGSA-N | Organic compounds | Lipids and lipid-like molecules | Steroids and steroid derivatives | Stigmastanes and derivatives | Stigmastanes and derivatives |
| **10630** | 24-Methylcholest-5,22-dien-3-ol; (3-beta,22E,24R)-form | OILXMJHPFNGGTO-BQYQJAHWSA-N | Organic compounds | Lipids and lipid-like molecules | Steroids and steroid derivatives | Ergostane steroids | Ergosterols and derivatives |
| **1774** | beta-Sitosterol | KZJWDPNRJALLNS-UHFFFAOYSA-N | Organic compounds | Lipids and lipid-like molecules | Steroids and steroid derivatives | Stigmastanes and derivatives | Stigmastanes and derivatives |
| **10631** | 24-Ethyl-22-trans-dehydrocholesterol; (3-beta,22E,24R)-form | HCXVJBMSMIARIN-CMDGGOBGSA-N | Organic compounds | Lipids and lipid-like molecules | Steroids and steroid derivatives | Stigmastanes and derivatives | Stigmastanes and derivatives |
| **1776** | Brassicasterol | OILXMJHPFNGGTO-BQYQJAHWSA-N | Organic compounds | Lipids and lipid-like molecules | Steroids and steroid derivatives | Ergostane steroids | Ergosterols and derivatives |
| **1777** | Cholesterol | HVYWMOMLDIMFJA-UHFFFAOYSA-N | Organic compounds | Lipids and lipid-like molecules | Steroids and steroid derivatives | Cholestane steroids | Cholesterols and derivatives |
| **2550** | Campest-5-en-7-one-3-beta-ol | LTLKHSBYMNKWPF-UHFFFAOYSA-N | Organic compounds | Lipids and lipid-like molecules | Steroids and steroid derivatives | Ergostane steroids | Ergosterols and derivatives |

**Table S10.** List of chemicals which are co-targeted on S5R1 and AR.

| **Chemical ID** | **Chemical Name** | **inchikey** | **Chemical Class** | | | | |
| --- | --- | --- | --- | --- | --- | --- | --- |
|  |  |  | **Kingdom** | **Superclass** | **Class** | **Subclass** | **Direct_parent** |
| **1984** | Dihydro-epideoxyarteannuin B | SXDUGGRDNCRRHY-UHFFFAOYSA-N | Organic compounds | Lipids and lipid-like molecules | Prenol lipids | Terpene lactones | Germacranolides and derivatives |
| **9607** | 4-Hydroxy-4-methyl-5-dodecanylcyclopenten-2-one | HWMQXCKSSOGAEL-UHFFFAOYSA-N | Organic compounds | Organic oxygen compounds | Organooxygen compounds | Alcohols and polyols | Tertiary alcohols |
| **3815** | 19(4-3)Abeo-12,14,15-trihydroxy-11-methoxyabiet-4(18),8,11,13-Tetraen-7-one | NHLQPIOUCQBSFL-UHFFFAOYSA-N | Organic compounds | Lipids and lipid-like molecules | Prenol lipids | Diterpenoids | Diterpenoids |
| **9612** | Kusunol | MQWIFDHBNGIVPO-UHFFFAOYSA-N | Organic compounds | Lipids and lipid-like molecules | Prenol lipids | Sesquiterpenoids | Eremophilane, 8,9-secoeremophilane and furoeremophilane sesquiterpenoids |
| **1847** | (-)-alpha-Copaene | VLXDPFLIRFYIME-UHFFFAOYSA-N | Organic compounds | Lipids and lipid-like molecules | Prenol lipids | Sesquiterpenoids | Sesquiterpenoids |
| **10638** | Stigmasta-4,6-dien-3-one | KEAZWUZFBSXOMV-UHFFFAOYSA-N | Organic compounds | Lipids and lipid-like molecules | Steroids and steroid derivatives | Stigmastanes and derivatives | Stigmastanes and derivatives |
| **2253** | epsilon-Muurolene | NOLWRMQDWRAODO-UHFFFAOYSA-N | Organic compounds | Lipids and lipid-like molecules | Prenol lipids | Sesquiterpenoids | Sesquiterpenoids |
| **2001** | Stigmastan-3-one (5-alpha) | BVVFRHKBULZQCQ-UHFFFAOYSA-N | Organic compounds | Lipids and lipid-like molecules | Steroids and steroid derivatives | Stigmastanes and derivatives | Stigmastanes and derivatives |
| **11158** | Dehydroabietinal | YCLCHPWRGSDZKL-UHFFFAOYSA-N | Organic compounds | Lipids and lipid-like molecules | Prenol lipids | Diterpenoids | Diterpenoids |
| **9654** | (-)-Ylangene | VLXDPFLIRFYIME-UHFFFAOYSA-N | Organic compounds | Lipids and lipid-like molecules | Prenol lipids | Sesquiterpenoids | Sesquiterpenoids |

**Table S11.** List of chemicals which are co-targeted on S5R2 and AR.

|  |  |  | **Chemical Class** | | | | |
| --- | --- | --- | --- | --- | --- | --- | --- |
| **Chemical ID** | **Chemical Name** | **inchikey** | **Kingdom** | **Superclass** | **Class** | **Subclass** | **Direct_parent** |
| **12777** | Zerumbone | GIHNTRQPEMKFKO-SKTNYSRSSA-N | Organic compounds | Lipids and lipid-like molecules | Prenol lipids | Sesquiterpenoids | Sesquiterpenoids |
| **9612** | Kusunol | MQWIFDHBNGIVPO-UHFFFAOYSA-N | Organic compounds | Lipids and lipid-like molecules | Prenol lipids | Sesquiterpenoids | Eremophilane, 8,9-secoeremophilane and furoeremophilane sesquiterpenoids |
| **1847** | (-)-alpha-Copaene | VLXDPFLIRFYIME-UHFFFAOYSA-N | Organic compounds | Lipids and lipid-like molecules | Prenol lipids | Sesquiterpenoids | Sesquiterpenoids |
| **10638** | Stigmasta-4,6-dien-3-one | KEAZWUZFBSXOMV-UHFFFAOYSA-N | Organic compounds | Lipids and lipid-like molecules | Steroids and steroid derivatives | Stigmastanes and derivatives | Stigmastanes and derivatives |
| **2001** | Stigmastan-3-one (5-alpha) | BVVFRHKBULZQCQ-UHFFFAOYSA-N | Organic compounds | Lipids and lipid-like molecules | Steroids and steroid derivatives | Stigmastanes and derivatives | Stigmastanes and derivatives |
| **11158** | Dehydroabietinal | YCLCHPWRGSDZKL-UHFFFAOYSA-N | Organic compounds | Lipids and lipid-like molecules | Prenol lipids | Diterpenoids | Diterpenoids |
| **9654** | (-)-Ylangene | VLXDPFLIRFYIME-UHFFFAOYSA-N | Organic compounds | Lipids and lipid-like molecules | Prenol lipids | Sesquiterpenoids | Sesquiterpenoids |
